# Understanding when and how knowledge graphs add value to EHR-based clinical prediction across Parkinson’s disease, multiple sclerosis, and chronic pain

**DOI:** 10.64898/2026.10.04.756605

**Authors:** Aishwarya Pathak, Oleg Stroganov, Tracy Ballinger, Tania Khasanova

**Affiliations:** Rancho Biosciences, Rancho Santa Fe, CA 92067, USA

## Abstract

Biomedical knowledge graphs (KGs) have emerged as a promising approach for incorporating biological knowledge into machine-learning models trained on electronic health record (EHR) data. However, their large scale and heterogeneous structure impose substantial computational cost, and it remains unclear whether their benefits generalize across diseases or depend on how the graph is constructed. We systematically evaluated KG integration across three diseases spanning distinct biological and clinical characteristics: multiple sclerosis, Parkinson’s disease, and chronic pain. For each disease we compared five modelling strategies, spanning EHR-only features and graph diffusion over concept-level, cohort-derived, disease-anchored, and full SPOKE (Scalable Precision Medicine Open Knowledge Engine) subgraphs, asking whether graph construction affects predictive performance and computational cost, whether KG integration improves classification over EHR-only models, and what the graph does to model behavior. Cohort-derived subgraphs, comprising roughly 45% of SPOKE’s nodes and one-fifth of its edges, matched or modestly exceeded the full graph’s performance while substantially reducing computational cost in all three diseases. The predictive benefit of KG integration, however, was strongly disease-dependent: it improved classification in Parkinson’s disease, had no measurable effect in multiple sclerosis, and reduced performance in chronic pain. These differences were not explained by the amount of biological information incorporated: the disease with the largest shift toward molecular features showed no improvement, the sole beneficiary showed only a modest shift, and the disease with no molecular shift was penalized. Mechanistically, graph diffusion redistributed each disease’s existing signal across many correlated features rather than adding independent biological signal, so that any gain depended on whether it added information beyond the EHR. These findings show that cohort-derived subgraphs offer a computationally efficient alternative to full-scale knowledge graphs, and that KG integration functions less as a universal accuracy booster than as a data- and disease-dependent transformation of the patient representation.

## Introduction

### Knowledge Graphs for EHR-Based Prediction

Every medical encounter generates a vast amount of clinical data- a diagnosis recorded, a medication prescribed, a laboratory result returned, which leaves a structured trace in the electronic health record (EHR). These data provide a rich resource for studying patient health, including the identification of patterns that predict future diseases [1]. Machine learning models trained on EHR data have begun to realize this potential, demonstrating meaningful predictive power across a range of clinical conditions [2, 3]. Yet EHR data alone has an important limitation: it captures *what* was observed about a patient, but not *why* those observations are biologically connected. A patient may carry a diagnosis, take a medication, and harbor a genetic variant that is all deeply related through shared molecular mechanisms, but the EHR encodes them as separate entries with no explicit link between them.

Biomedical knowledge graphs offer a way to address this limitation. At their core, Knowledge Graphs (KGs) are structured representations of entities and the relationships between them. Resources such as SPOKE (Scalable Precision Medicine Open Knowledge Engine)[4], HetioNet [5], and PrimeKG [6] encode biological reality in structured form: connecting genes to pathways, compounds to targets, diseases to molecular mechanisms, derived from curated experimental evidence rather than clinical documentation. The intuitive hypothesis is that bridging these two representations should improve disease prediction: grounding EHR-derived patient profiles in biological context should allow models to move beyond administrative coding patterns toward mechanistically meaningful features.

A key methodological challenge is how to derive compact, informative representations from knowledge graphs for use in downstream prediction tasks. Diffusion-based methods have proven particularly well-suited to this purpose, propagating information across graph topology from seed nodes corresponding to a patient’s observed EHR concepts and yielding embeddings that capture both direct and higher-order biological relationships [7]. Nelson et al. 2022 demonstrated this approach by embedding patient EHR concepts into SPOKE using a personalized PageRank algorithm to generate patient-level graph embeddings termed SPOKEsigs [8]. Applied to Multiple Sclerosis (MS) prediction, the model demonstrated strong predictive performance and highlighted biologically relevant features consistent with known MS mechanisms [8], establishing a proof of concept that diffusing EHR-derived signals through a biomedical knowledge graph can yield informative patient representations for clinical prediction.

### Limitations of Knowledge Graphs

Despite this promise, several fundamental questions remain unanswered. First, it is unclear whether knowledge graph integration genuinely improves prediction over standard machine learning applied to raw EHR features, or whether the apparent benefit reflects the strength of the EHR signal itself. Second, universal biomedical knowledge graphs such as SPOKE [4] present substantial practical challenges: operating at full scale imposes significant computational burden, and the breadth that makes universal graphs valuable as general-purpose resources may limit their effectiveness in focused clinical applications; diffusion across the full graph risks diluting disease-relevant signal with contributions from unrelated biological regions. Third, and most fundamentally, the mechanism by which knowledge graph integration influences model behavior remains poorly characterized: does the graph genuinely shift model reasoning toward biological mechanisms, or does it simply repackage the same clinical signal in a different representation?

One natural baseline for managing this computational burden is random subsampling of graph nodes. Although practical, this strategy is agnostic to biological and clinical context. Node selection does not account for disease relevance or patient population, and subsampled graphs may omit informative biological structure while preserving irrelevant components. More targeted graph construction strategies may yield both computational efficiency and improved biological fidelity, but this has not been systematically evaluated. Critically, all prior work has evaluated knowledge graph integration in single-disease settings, making it impossible to determine whether observed benefits are general properties of graph integration or specific to the disease and EHR context studied [8, 9]. Whether KG integration helps, is neutral, or actively hurts prediction may depend on properties of the disease itself: its biological specificity, its representation in the knowledge graph, and the sufficiency of its EHR footprint, but this has never been tested across a systematic multi-disease comparison.

In this study, we address these gaps through a systematic evaluation of KG integration across three diseases representing a deliberate spectrum of biological and clinical characteristics: Parkinson’s disease (PD), a well-characterized neurodegenerative disease with a specific drug treatment footprint and rich SPOKE representation [4]; multiple sclerosis (MS), a well-characterized autoimmune disease with strong EHR signal and good SPOKE representation; and Chronic Pain, a heterogeneous syndrome with strong but non-specific EHR signal and sparse SPOKE representation. We evaluate multiple graph construction strategies per disease: from full SPOKE traversal to maximally pruned patient-anchored subgraphs.

We explore three broad questions: (i) whether the way the knowledge graph is constructed and scaled affects predictive performance and computational cost; (ii) whether integrating the knowledge graph improves prediction over using the EHR data alone, and whether this depends on the disease; and (iii) what the knowledge graph is doing to the model mechanistically. Together, these questions address a central challenge in the field: **whether integrating biomedical knowledge graphs with EHR data improves disease classification and, if so, under what conditions, how the graph should be incorporated, and what information it contributes beyond the EHR alone?**

We show that cohort-driven subgraphs consisting of approximately 45% of SPOKE’s nodes and roughly one-fifth of its edges, match or modestly exceed full-graph performance across all three diseases at a fraction of the computational cost. We further show that the accuracy benefit of KG integration is strongly disease-dependent, ranging from improvement in Parkinson’s disease to no change in multiple sclerosis to deterioration in chronic pain, and, critically, that this benefit is not a function of how much biological signal the graph introduces: the disease with the largest molecular shift gained nothing, whereas the disease with a far smaller shift was the sole beneficiary. Feature-level analysis indicates that the graph redistributes each disease’s existing signal across many correlated features, so that any gain depends on whether diffusion adds information beyond the EHR. Knowledge-graph integration thus functions less as a universal accuracy booster than as a biological translation layer, whose value extends beyond discrimination to the interpretability and biological grounding of clinical predictions, even where it does not improve, or occasionally reduces predictive accuracy.

## Results

### A systematic comparison of graph construction strategies across diverse disease cohorts

Patient-level knowledge graph diffusion was originally introduced as a proof of concept in a single disease, where electronic health record (EHR) concepts were embedded into the complete SPOKE knowledge graph using personalized PageRank to generate patient-specific signatures (*SPOKEsigs*) [8] (illustrated in Fig. 1a). Although this approach demonstrated the value of integrating EHR data with biomedical knowledge, it did not address whether diffusion across the complete graph is necessary for prediction or whether comparable performance can be achieved using only the graph regions most relevant to a given patient population. To address this question systematically, we developed a comparative evaluation framework that varied both disease context and graph scope, enabling the contribution of graph construction to be assessed independently across diseases rather than within a single proof-of-concept cohort (Fig. 1).

**Figure 1.**
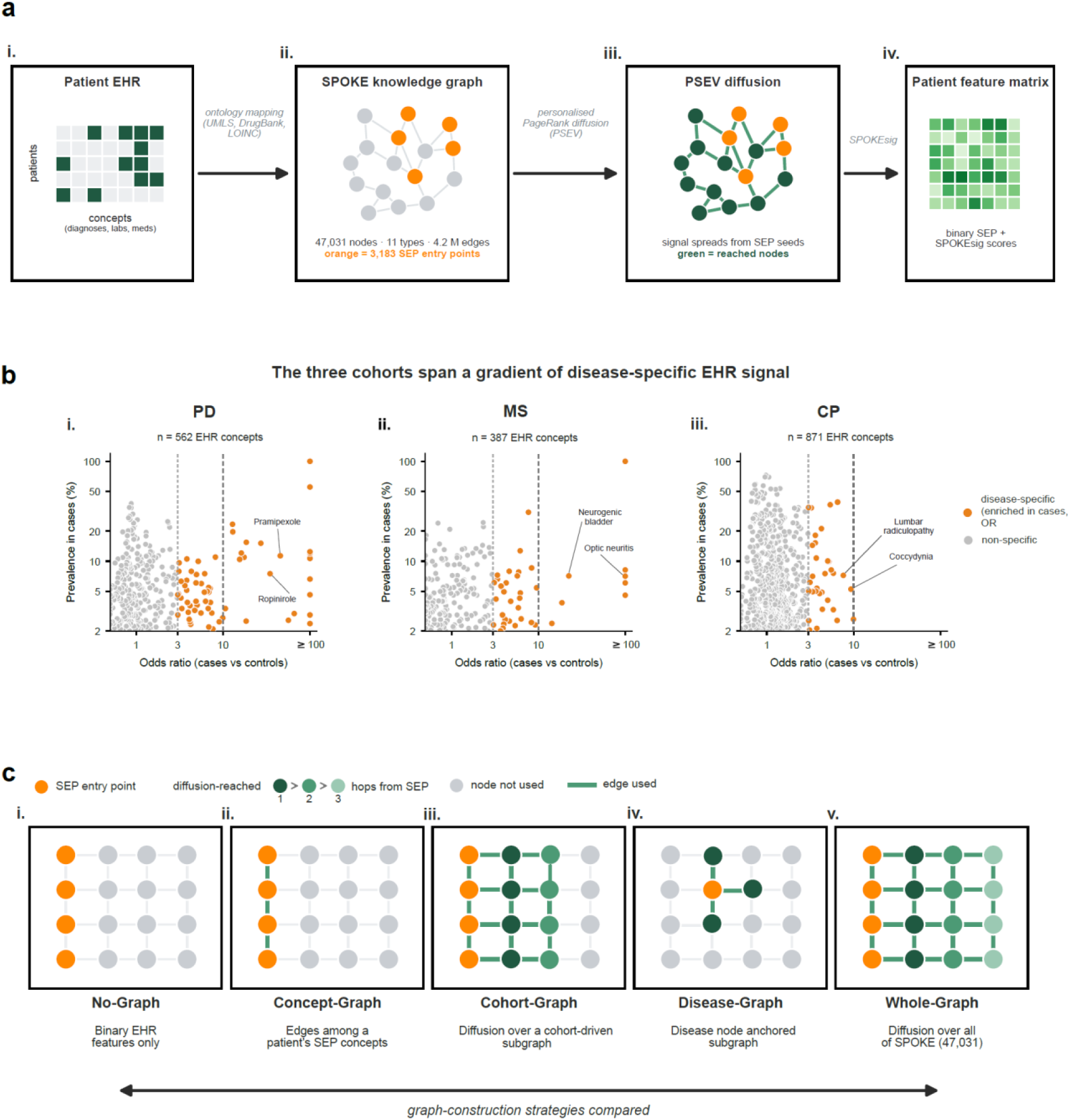
A systematic framework for comparing knowledge-graph construction strategies across diverse disease cohorts. (a) Each patient’s electronic health record (EHR is represented as a binary patient × concept matrix and mapped, via standard ontologies (UMLS, DrugBank, LOINC), onto the SPOKE knowledge graph (47,031 nodes across 11 types, ∼4.2 M edges). The 3,183 mapped concepts are the SPOKE entry points (SEPs; orange). Personalized-PageRank (PSEV) diffusion propagates each patient’s signal from their SEP seeds across the graph (green = reached nodes, green edges = edges traversed); the resulting SPOKEsig scores, with the binary SEP features, form the patient feature matrix used for classification. (b) The three cohorts span a gradient of disease-specific EHR signal. Each dot is one EHR concept present in ≥2% of a cohort’s cases (n = 562 PD, 387 MS, 871 CP), positioned by specificity (x: odds ratio, cases vs matched controls, Fisher’s exact, log) and strength (y: prevalence in cases, log). Concepts enriched in cases (OR ≥ 3, FDR-corrected) are orange (disease-specific; 89 PD, 48 MS, 38 CP), others gray; dashed lines mark OR = 3 and OR = 10 (OR capped at 100 for display). Representative disease-specific concepts are labelled (PD: Pramipexole, Ropinirole; MS: Optic neuritis, Neurogenic bladder; CP: Lumbar radiculopathy, Coccydynia). PD and MS carry strong, specific signal where many concepts exceed OR = 10, whereas CP has strong but non-specific signal. (c) Five graph-construction strategies illustrated, on a shared SPOKE lattice with color on a gray base (orange = SEP entry point; green = diffusion-reached, shaded by hop 1-3; gray = present but unused; green line = edge traversed). (i) No-Graph: binary SEP features only. (ii) Concept-Graph: edges among a patient’s own SEP concepts only. (iii) Cohort-Graph: diffusion over a cohort-derived subgraph - all cohort SEPs plus their one-hop neighbors, hubs removed. (iv) Disease-Graph: diffusion over a disease-centered subgraph - the disease node plus its one-hop neighborhood (322 nodes for MS); anchored on the disease rather than patient concepts, only ∼3% of SEPs fall inside. v) Whole-Graph: diffusion over all of SPOKE, identical to (a).

To capture a range of clinical and biological settings, we selected three balanced case-control cohorts that differ systematically in disease specificity, EHR signal strength and knowledge-graph representation (Fig. 1b). For every EHR concept present in at least 2% of a cohort’s cases, we quantified its strength (prevalence in cases) and its specificity (enrichment in cases relative to matched controls; odds ratio, Fisher’s exact test). Parkinson’s disease (PD; n = 4,298) is a neurodegenerative disorder with an even more distinctive, pharmacologically driven profile and similarly rich graph representation: 89 of 562 concepts were disease-specific, dominated by dopaminergic therapies (e.g., pramipexole, ropinirole) at very high odds ratios (Fig. 1b.i). Multiple sclerosis (MS; n = 4,698) is a well-characterized autoimmune disease with strong, highly specific EHR signal and extensive representation within SPOKE: of 387 frequent concepts, 48 were disease-specific (OR ≥ 3, FDR-significant), many exceeding OR = 10 and including hallmark features such as optic neuritis and neurogenic bladder (Fig. 1b.ii). In contrast, chronic pain (CP; n = 23,748) is a heterogeneous syndrome with diffuse clinical manifestations and comparatively sparse graph representation: although it comprised the most frequent concepts (871), only 38 were disease-specific, reflecting strong but non-specific signal in which characteristic concepts occur at similar rates in cases and matched controls (Fig. 1b.iii). Together, these cohorts span a spectrum - from specific and graph-rich (MS, PD) to non-specific and graph-sparse (CP), across which the value of knowledge-graph diffusion may reasonably differ.

Within each cohort, we evaluated five modelling strategies that progressively increased the incorporation of graph information (illustrated in Fig. 1c). The **“No-Graph”** model served as a graph-free baseline, training only on binary EHR (SEP) features (Fig. 1c.i). Four additional approaches generated patient embeddings through personalized PageRank diffusion but differed in the graph over which diffusion was performed. The **“Concept-Graph”** consisted of the knowledge graph subgraph connecting concepts present within an individual patient’s EHR, providing a minimal graph representation without knowledge propagation (Fig. 1c.ii). The **“Cohort-Graph”** restricted diffusion to a one-hop expansion surrounding concepts observed within the study cohort, producing subgraphs of approximately 21,200 nodes (Fig. 1c.iii). The **“Disease-Graph”** represented the most aggressive reduction, limiting diffusion to compact disease-centered subgraphs comprising 322-798 nodes derived from disease seed concepts (Fig. 1c.iv). Finally, the **“Whole-Graph”** reproduced the original SPOKEsigs framework by diffusing across all 47,031 SPOKE nodes (Fig. 1c.v). Together, these comparisons isolate the contribution of graph scope independently of the underlying learning algorithm, providing a systematic framework for evaluating knowledge-graph construction across diseases.

### Cohort-focused graph construction matches whole-graph performance at a fraction of the computational cost

To determine how graph construction influenced predictive performance, we compared the three diffusion-based strategies that differed only in the graph over which personalized PageRank was performed. We first asked whether diffusion across the complete SPOKE graph (Whole-Graph) provided any advantage over the more focused Cohort-Graph. The logic was simple: if restricting diffusion to a smaller cohort- or disease-derived subgraph discarded useful biology, the reduced models should underperform the Whole-Graph.

Model performance was evaluated using AUROC across 25 independent model-building instances (each a single stratified train/test split), with pairwise comparisons assessed using paired t-tests with Bonferroni correction and Cohen’s d. Because some diseases contained highly prevalent, disease-specific EHR concepts that enabled near-trivial classification (Fig. 1b), these disease-defining features were excluded from all subsequent analyses (see Methods, *Leakage Removal).* In every case they were direct markers of the disease: for Parkinson’s disease- the PD diagnosis and its defining medications, such as L-DOPA (levodopa, the primary dopamine-replacement therapy, present in 55% of cases), rasagiline and entacapone; for multiple sclerosis (MS- the MS diagnosis itself, the MS-associated drugs fingolimod and modafinil, and optic neuritis (a hallmark presenting symptom); and for chronic pain, the symptom concepts that define the syndrome (back pain, low back pain and loin pain). Removing these features ensured that model performance reflected the contribution of knowledge-graph integration rather than obvious diagnostic or treatment markers (full list with prevalences in Supplementary Table 2). Because the choice of which features to exclude could in principle influence these comparisons, we repeated the entire pipeline under a less-stringent removal scheme that retained these disease-defining features; every conclusion reported here and below was unchanged, with only modest shifts in absolute AUROC (see Methods, *Sensitivity to leakage-removal stringency*; Supplementary Fig. 2).

Across all three cohorts, the Whole-Graph did not outperform the Cohort-Graph. If anything, the Graph was marginally higher in PD (d = +0.976, P < 0.001), MS (d = +0.791, P = 0.0059) and CP (d = +3.369, P < 0.001), but with near-identical mean AUROCs (PD 0.950 vs 0.949; MS 0.945 vs 0.944; CP 0.808 vs 0.805). Figure 2a-i plots these as the per-instance difference (Cohort-Graph − Whole-Graph) across the 25 matched splits; in every cohort the differences fell within |Δ| < 0.005 of zero. They reached statistical significance because the tightly paired design and low across-split variance make even trivial offsets detectable, but in absolute terms they were negligible and, where non-zero, favored the smaller Cohort-Graph. The Cohort-Graph is therefore predictively equivalent to the Whole-Graph, expanding diffusion to the complete SPOKE graph conferred no benefit.

**Figure 2.**
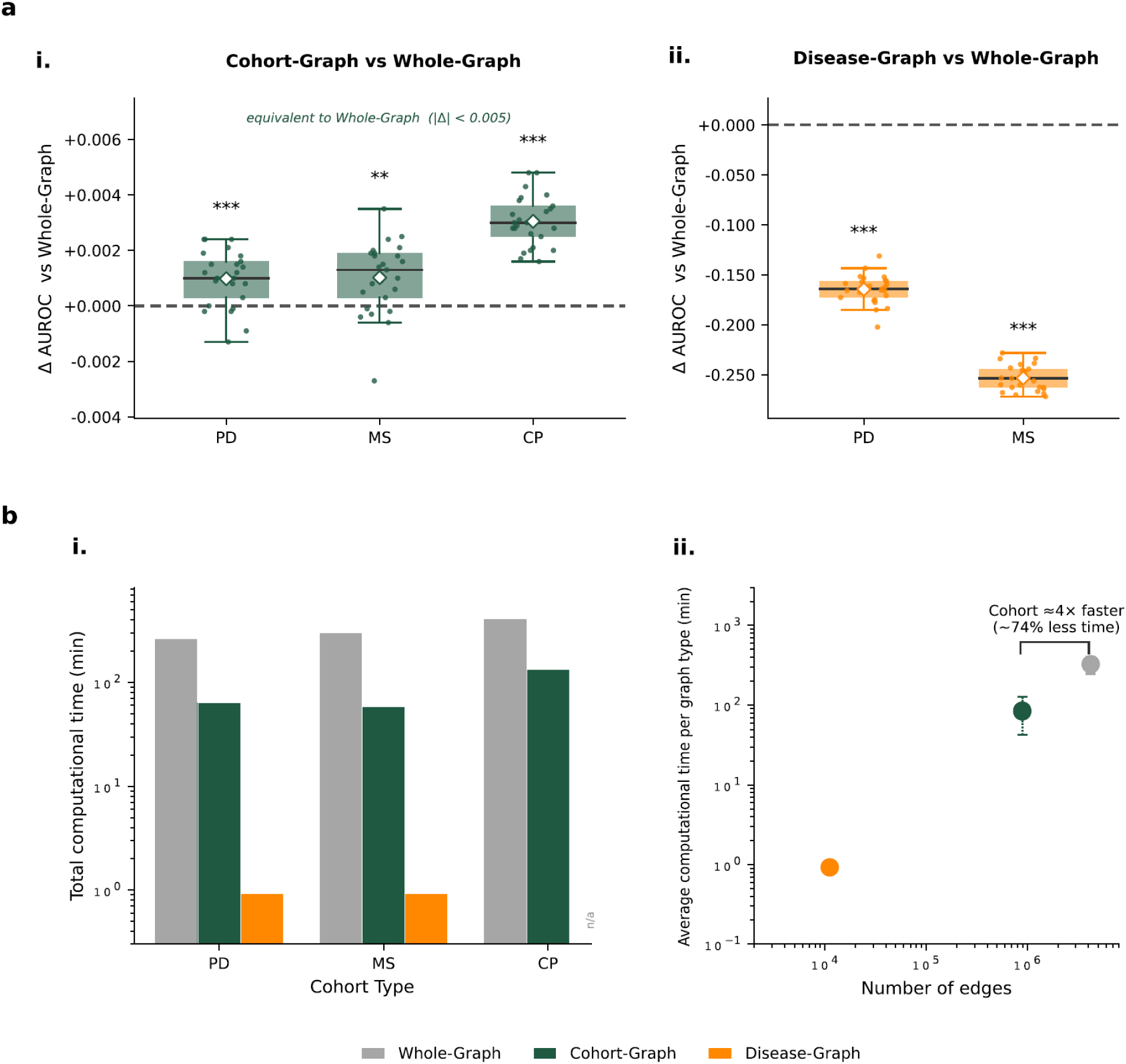
Graph construction strategy shapes predictive performance and computational cost. (a) Predictive performance relative to Whole-Graph, per model-building instance. For each of 25 independent build instances, paired difference in test AUROC between a reduced-graph strategy and the Whole-Graph model were evaluated on the same split. (i) Cohort-Graph minus Whole-Graph across all three cohorts (PD, MS, CP): differences cluster tightly around zero (|Δ| < 0.005 in every cohort) - Cohort-Graph is predictively equivalent to, and marginally exceeds, Whole-Graph. (ii) Disease-Graph minus Whole-Graph (PD and MS only; chronic pain has no disease-anchored subgraph): performance collapses (MS ≈ −0.25, PD ≈ −0.16 AUROC). Boxes show median and interquartile range, points the 25 per-instance deltas, and the white diamond the mean; the dashed line marks zero. Significance from two-tailed paired t-tests, Bonferroni-corrected (** q < 0.01, *** q < 0.001). (b) Computational cost. (i) Total computational time (personalized-PageRank diffusion plus random-forest training) for each graph type within each cohort (PD, MS, CP; log-scaled y-axis). Whole-Graph (gray) is the most expensive strategy in every cohort, Cohort-Graph (green) is 3.1- to 5.2-fold cheaper, and Disease-Graph (orange) is trivially cheap (∼0.9 min); Disease-Graph was not constructed for chronic pain (marked "n/a"). (ii) The same total time, averaged across the three cohorts for each graph type, plotted against the number of edges in the graph diffused over (log-log). Points are the per-graph-type mean across cohorts and error bars the standard deviation.

While the two strategies were comparable in accuracy, they differed markedly in computational cost. Using identical hardware and parameters (Methods, *Computational Parameters*), we quantified each strategy’s total cost as graph diffusion (personalized PageRank) plus model training and evaluation across the 25 independent train/test builds (Fig. 2b). The same ordering held in all three cohorts: the Whole-Graph was the most computationally intensive and the more focused graphs progressively less demanding. For the Whole-Graph, runtime was dominated by diffusion over the full SPOKE graph, requiring 262.2 min (∼4.4 h) in PD, 299.2 min (∼5.0 h) in MS and 412.4 min (∼6.9 h) in CP. In contrast, the Cohort-Graph matched Whole-Graph accuracy at a fraction of the cost - 63.1, 58.0 and 132.5 min (∼1.1, 1.0 and 2.2 h) in PD, MS and CP respectively - a 3.1- to 5.2-fold reduction in runtime, achieved by diffusing over a hub-pruned subgraph that retained only ∼45% of the nodes and ∼21% of the edges of the full SPOKE graph (∼21,000 vs 47,031 nodes; ∼0.89M vs 4.2M edges). The Disease-Graph, evaluated only in MS and PD, was the least expensive to construct (∼0.9 min in both cohorts), but this reduction came at the expense of predictive performance, as described in the next section.

Runtime scaled with graph size (Fig. 2b-ii): the near-fivefold reduction in edges achieved by the Cohort-Graph accounts for its roughly fourfold lower diffusion cost. Absolute runtimes were highest in chronic pain for every strategy, reflecting its substantially larger cohort (n = 23,748) and correspondingly greater training cost; nevertheless, the relative computational advantage of the focused Cohort-Graph was preserved across all cohorts.

### Disease-centered graph construction substantially impairs predictive performance

Having established that diffusion could be restricted to a cohort-derived subgraph without loss of predictive performance, we next examined whether a smaller, disease-centered subgraph was sufficient. We found that restricting diffusion to the Disease-Graph substantially impaired classification (Fig. 2a.ii). The Disease-Graph yielded mean AUROCs of 0.7846 in PD and 0.6905 in MS, compared with ∼0.95 (PD) and ∼0.94 (MS) for both the Whole-Graph and Cohort-Graph strategies (all paired comparisons P < 0.001; Cohen’s |d| = 11.4 and 11.6 in PD, and 19.9 and 20.4 in MS, versus the Whole-Graph and Cohort-Graph respectively). This reduction reflected the graph construction itself: the disease-centered subgraph retained only ∼2-3% of each cohort’s SPOKE entry points (78 in PD and 86 in MS), excluding most patient-specific information before diffusion was performed. The strategy could not be applied to chronic pain because of how the condition is represented within SPOKE: rather than a biological disease node, chronic pain is encoded as terminal ’Symptom’ and ’Side-effect’ annotation nodes with few biological connections, yielding a seed subgraph of only 13 nodes (∼11 edges) - insufficient graph structure for meaningful personalized PageRank diffusion.

Together, these results establish graph scope as a design variable in its own right: predictive performance did not increase monotonically with graph size, and the most expensive option was not the best.

### Knowledge-graph integration exerts disease-dependent effects on predictive performance

Having established that the Cohort-Graph reproduces the predictive performance of the full Whole-Graph while using under half of SPOKE (Fig. 2), we adopted it as the reference graph-based strategy and asked how two lower-scope strategies compared: the No-Graph baseline (a random forest on the binary SEP features, with no graph diffusion) and the Concept-Graph (diffusion restricted to edges among a patient’s own SEP concepts, without expansion into the surrounding graph). Each strategy was compared to the Cohort-Graph within each cohort across 25 independent train/test builds (two-tailed paired t-tests, Bonferroni-corrected, Cohen’s d; Fig. 3 plots per-instance differences relative to the Cohort-Graph).

**Figure 3.**
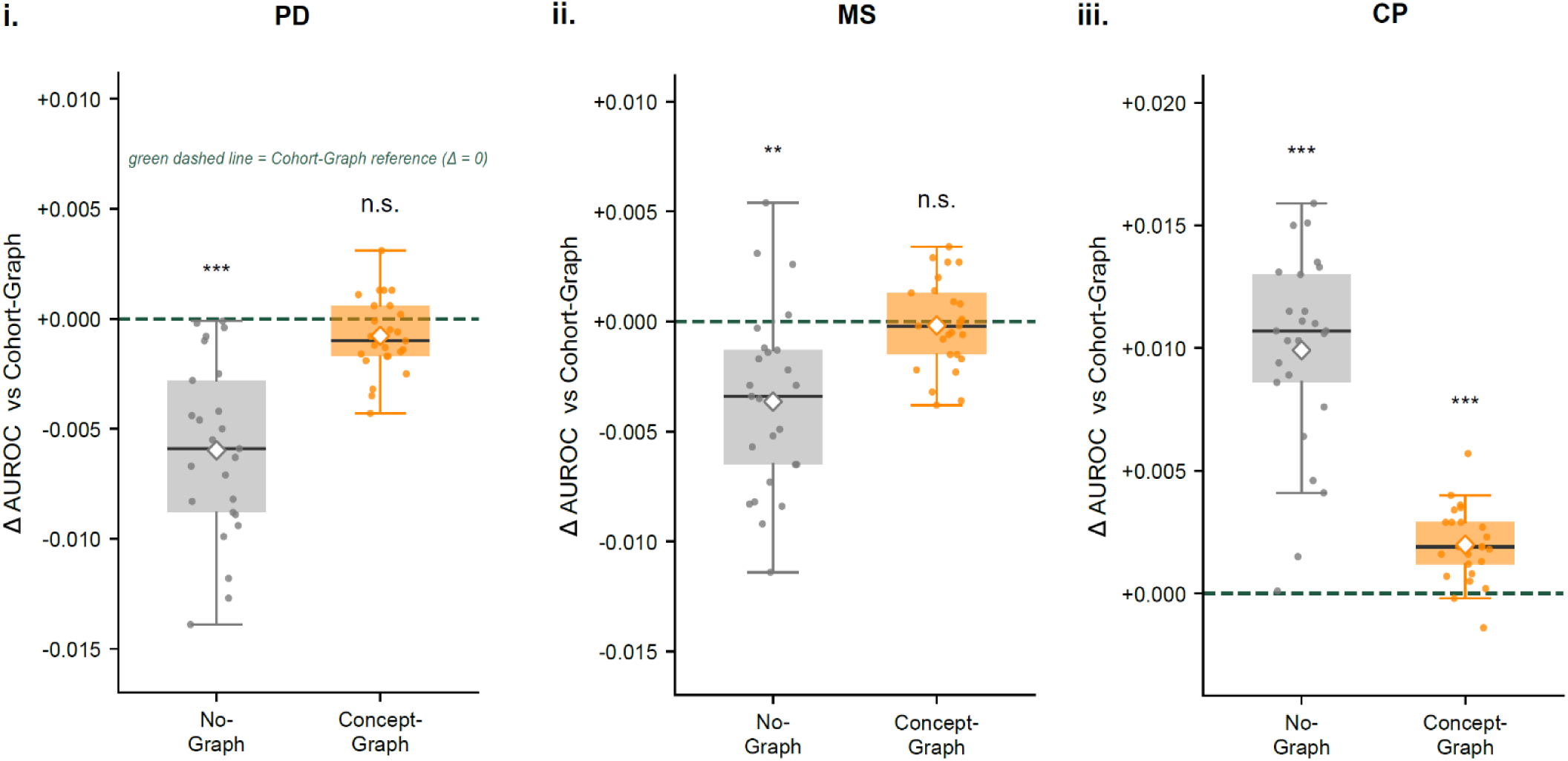
The predictive value of knowledge-graph integration is disease-dependent. Per-instance AUROC differences relative to the Cohort-Graph model (green dashed line, Δ = 0) for the two lower-integration strategies - No-Graph (binary EHR/SEP features, no diffusion; gray) and Concept-Graph (SEP plus intra-SEP edges; orange). Panels: (i) Parkinson’s Disease (PD), (ii) Multiple Sclerosis (MS), (iii) chronic pain (CP). Boxes show median and interquartile range, points the 25 per-instance deltas, and the white diamond the mean; significance is from two-tailed paired t-tests against the Cohort-Graph, Bonferroni-corrected (** q < 0.01, *** q < 0.001; n.s., not significant). In PD and MS, the No-Graph model performed significantly worse than the Cohort-Graph (PD Δ ≈ −0.006; MS Δ ≈ −0.004), while the Concept-Graph was statistically indistinguishable from it (n.s.)- graph diffusion over the cohort-derived subgraph therefore adds predictive value beyond raw EHR features. In CP the pattern reversed: both the No-Graph (Δ ≈ +0.010) and the Concept-Graph (Δ ≈ +0.002) performed significantly better than the Cohort-Graph, indicating that graph integration degraded rather than improved prediction.

The effect of graph integration was strongly disease-dependent (Fig. 3). In PD and MS, discarding graph diffusion degraded performance: the No-Graph baseline fell significantly below the Cohort-Graph (PD ΔAUROC = −0.006, d = 1.50, p < 0.001; MS Δ = −0.004, d = 0.88, p = 0.002). The Concept-Graph, however, was statistically indistinguishable from the Cohort-Graph in both cohorts (PD Δ = −0.001, p = 0.34; MS Δ ≈ 0.000, p = 1.0), indicating that the edge structure among a patient’s own concepts already captured the available graph signal, and that one-hop expansion into the cohort subgraph added nothing further. In CP the pattern inverted: every graph-based strategy performed worse than none, and performance declined monotonically as more graph structure was incorporated - the No-Graph baseline significantly exceeded the Cohort-Graph (Δ = +0.010, d = 2.45, p < 0.001), with the Concept-Graph sitting between the two but still significantly above the Cohort-Graph (Δ = +0.002, d = 1.32, p < 0.001). Although the absolute differences were small, they were highly consistent across the 25 paired builds and reproducible in direction within each cohort. Notably, the magnitude of the harm in chronic pain (No-Graph vs Cohort, d = 2.45) exceeded the benefit of the graph in Parkinson’s disease (d = 1.50), so integration was neither uniform nor consistently beneficial across the three cohorts.

### Graph diffusion distributes predictive signal across correlated features

We showed that knowledge-graph integration altered predictive performance in a disease-dependent manner, ranging from improvement to deterioration across the three cohorts. However, these results do not explain how graph integration changes model behavior to produce these contrasting outcomes. We therefore asked whether graph integration systematically shifts the model toward biologically informed features, or simply redistributes importance among existing clinical features, and whether such shifts are associated with improved prediction. To address these questions, we quantified permutation feature importance for each modelling strategy and disease cohort (Fig. 4).

**Figure 4.**
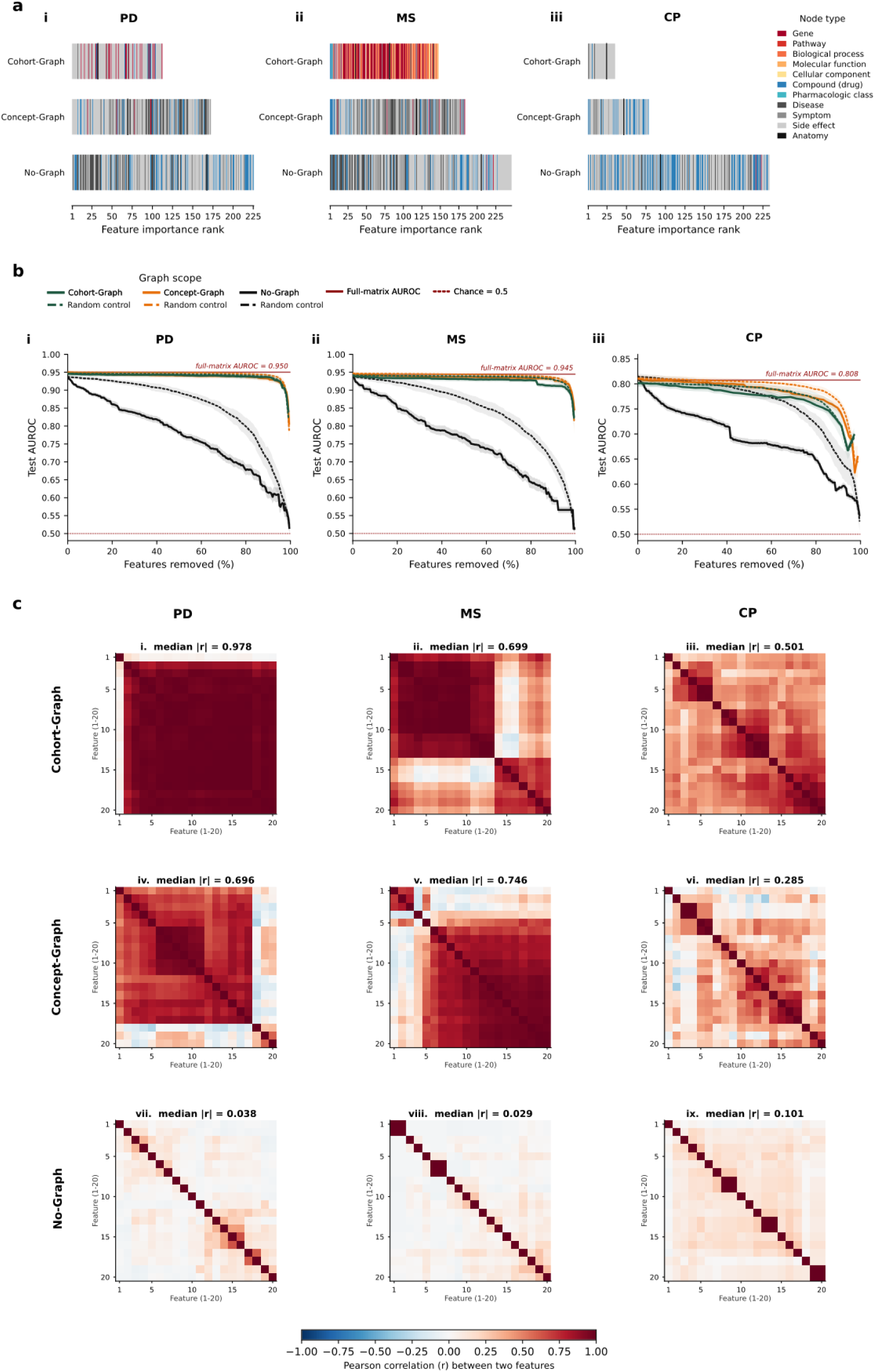
Wider graph scope yields more correlated, distributed feature importance. (a) Recruited features for each graph scope (rows, top to bottom: Cohort-Graph, Concept-Graph, No-Graph) and cohort (i, PD; ii, MS; iii, CP), arranged left- to-right by permutation-importance rank and colored by SPOKE node types. Strip length equals the total number of recruited features. (b) Feature ablation for each cohort (i, PD; ii, MS; iii, CP). Recruited features were removed cumulatively, either most-important-first (solid lines) or in random order (dotted lines; control), retraining the random forest at each step; lines show the mean and shaded bands ±1 standard deviation of test AUROC across all builds, for the three graph scopes (Cohort-Graph, green; Concept-Graph, orange; No-Graph, black). The dark-red solid line marks the Cohort-Graph full-matrix (all-features) AUROC; the dark-red dotted line marks chance (0.5). In the No-Graph models, importance-ordered removal falls steeply below the random-order control, whereas the Concept-Graph and Cohort-Graph curves remain near the full-matrix AUROC until nearly all features are removed, indicating that predictive signal is distributed across many features rather than concentrated in a few. (c) Pairwise Pearson correlation among the top 20 recruited features (by permutation importance) for each graph scope (rows: Cohort-Graph, i–iii; Concept-Graph, iv–vi; No-Graph, vii–ix) and cohort (columns: PD, MS, CP), with features ordered within each panel by hierarchical clustering. Color indicates the correlation coefficient r; the median absolute pairwise correlation (median |r|) is shown above each panel. Graph-derived features are far more correlated than No-Graph features, consistent with the flat ablation curves in (b).

Because random forests spread importance across many weak and correlated features, we quantified importance in two stages. First, to focus on the features each model actually used, we averaged every feature’s Gini importance- the impurity reduction it provided during training, across the 25 independent builds and retained those with mean Gini importance above 0.001 (the "recruited" set); this step identifies the features a model relied on but, being derived from the training data, does not measure their predictive value. We then ranked the recruited features by permutation importance, i.e., the mean decrease in held-out test AUROC across the 25 builds when a feature’s values were randomly shuffled, which quantifies each feature’s actual contribution to prediction; hereafter "importance" refers to this measure (Fig. 4a). Annotating each recruited feature by its SPOKE node type showed that graph integration reshaped both the size and the composition of the feature sets the models used. Without the graph, all three cohorts relied on large sets of raw EHR concepts dominated by clinical node types: in Parkinson’s disease the 225 recruited features comprised 106 side effects, 49 drugs, 47 diseases and 23 symptoms, with no genes (Fig. 4a.i), and multiple sclerosis was similar (the recruited 246 features comprised 126 side effects, 60 diseases, 43 drugs, 15 symptoms and 2 genes, Fig. 4a.ii). One-hop cohort-subgraph expansion changed these sets to differing degrees. In Parkinson’s disease the 112 recruited features remained largely clinical (77 side effects) but now included 24 genes (Fig. 4a.i); in multiple sclerosis the 147 recruited features were dominated by molecular entities (67 genes, 47 biological processes and 16 further pathway, molecular-function and cellular-component nodes, Fig. 4a.ii); and in chronic pain only 35 features were recruited, almost entirely symptom and side-effect nodes, with no molecular entities (Fig. 4a.iii).

A ranking alone, however, does not reveal whether these features carried independent or overlapping predictive signal. We therefore asked how many features were actually necessary for prediction, cumulatively removing recruited features: either most-important-first or, as a control, in random order, and retraining the model at each step (Fig. 4b). In the No-Graph models, importance-ordered removal produced a steep decline that fell well below the random-removal control, as expected when performance depends on a few non-redundant features (Fig. 4b). The Concept-Graph and Cohort-Graph models, however, behaved unexpectedly: removing features in importance order was scarcely more damaging than removing them at random, and AUROC remained near the full-model value until nearly all features had been removed. This indicated that in the graph models predictive signal was distributed across many features rather than concentrated in a few, with no single feature individually essential.

We hypothesized that this distributed importance reflected the way graph diffusion spreads signal: because propagation over the knowledge graph gives neighboring nodes similar values, a single disease-relevant signal is spread across many correlated features. Consistent with this hypothesis, the correlation structure of the recruited features differed sharply between strategies (Fig.4c). No-Graph features were essentially uncorrelated (median absolute pairwise correlation of 0.04, 0.03 and 0.10 in Parkinson’s disease, multiple sclerosis and chronic pain; Fig. 4c.vii-ix), whereas graph-derived features formed strongly correlated blocks, median |r| of 0.70, 0.75 and 0.29 at the Concept-Graph (Fig. 4c.iv-vi) and 0.98, 0.70 and 0.50 at the Cohort-Graph (Fig. 4c.i-iii), for the three cohorts. This explains the flat ablation curves for the graph models: because each disease’s signal is spread across many correlated features, removing any single feature-even the most important, has little effect, as the remaining correlated features still carry that signal.

## Discussion

Knowledge graphs are increasingly used to enrich electronic health record (EHR) representations with biological context, most prominently through the SPOKE framework, in which a patient’s EHR concepts are diffused over a large biomedical network to produce propagated entry vectors and patient signatures [4, 8, 10]. Prior applications have reported that SPOKE-embedded records can recognize prodromal disease and improve prediction in individual conditions such as multiple sclerosis [8] and Parkinson’s disease [9]. Building on that work, we asked three questions that single-disease studies leave unresolved: whether the way the graph is constructed matters, whether knowledge-graph integration helps at all and whether this varies by disease, and what the integration is doing to the model mechanistically. We addressed these across three cohorts spanning a range of biological specificity and representation within SPOKE: Parkinson’s disease, multiple sclerosis and chronic pain.

Graph construction proved to be a design choice in its own right. Predictive performance did not increase monotonically with graph size, and the most computationally expensive strategy was not the best. Diffusion over the complete SPOKE graph, as used in the original PSEV approach [10] offered no measurable predictive advantage over a far smaller cohort-derived graph, while the opposite extreme of a disease-centered subgraph discarded most patient-specific context and substantially impaired classification. A graph scoped to the concepts actually present in the patient population captured essentially all of the predictive signal available in SPOKE at a 3- to 5-fold lower computational cost (Fig. 2). Because network propagation is a diffusion process [7] whose cost scales with the number of edges traversed, pruning the graph translates directly into large runtime savings without a corresponding loss of signal. We therefore suggest a cohort-scoped graph as a practical default for knowledge-graph integration at scale, aligning graph scope to the data rather than maximizing or minimizing graph size.

Whether integration helped at all, however, depended on the disease. Across cohorts, its effect ranged from marginal improvement in PD to deterioration in CP (Fig. 3). This bidirectionality matters because the published SPOKE literature, like most graph-integration studies, has largely evaluated single diseases in which integration appeared beneficial [8, 9], and a single-disease design cannot separate a general property of the method from a characteristic of the disease under study. That integration can degrade prediction as readily as improve it is consistent with a broader pattern in the graph-representation-learning literature, where graph and graph-neural methods do not reliably outperform well-constructed tabular baselines and their advantage depends heavily on task and data [11, 12]. We frame the broader relationship cautiously: we observed that the impact of knowledge-graph integration differed across the three evaluated cohorts, which suggests that its utility may be influenced by disease and data characteristics rather than being assumed to provide a universal improvement. Whether that utility can be anticipated from disease or representational characteristics is a hypothesis that will require evaluation across a wider range of diseases and knowledge graphs.

The failure of integration in chronic pain also highlights how disease representation within the knowledge graph constrains what diffusion can achieve. Chronic pain is encoded in SPOKE largely as sparse terminal symptom and side-effect annotation nodes with few biological connections, so diffusion could not propagate beyond the clinical layer, and a disease-anchored subgraph could not be constructed at all. This is consistent with well-documented coverage and degree biases in biomedical knowledge graphs [13, 14], in which well-studied diseases and hub entities are densely connected while under-studied conditions remain sparsely represented. Knowledge-graph integration can therefore only contribute biologically where the underlying graph represents the disease with sufficient structure.

Examining the features that the models relied upon clarified a likely mechanism for these contrasting outcomes. Rather than adding a small set of features carrying information absent from the EHR, graph diffusion re-expressed each disease’s already-present predictive signal across many mutually correlated graph-derived features (Fig. 4). This reorganization is largely separate from whether prediction improves: because diffusion spreads signal that is in part already present in the EHR across many correlated graph-derived features, expanding the graph changes how information is represented more than how much information is available. Whether this helped, left performance unchanged, or harmed it then depended on whether diffusion contributed information beyond the EHR in a given disease. In PD it added modest signal, largely graph-derived clinical comorbidity features, and improved prediction; in multiple sclerosis the graph-derived features were largely correlated with an already-strong EHR signal and left performance unchanged, despite the largest shift toward molecular features; and in chronic pain, where the disease is sparsely represented in SPOKE, diffusion propagated non-specific clinical concepts and reduced performance. The predictive effect of integration therefore tracked whether the diffused representation added information beyond the EHR, not the amount of biology recruited or the size of the shift toward biological features. This distributed, correlated structure also carries an interpretive caution. Random forest importance estimates are diluted and biased when predictors are highly correlated [15, 16] and the correlation structure we observed is precisely the regime in which these rankings are least reliable, so the ranking of any single graph-derived feature should not be taken as evidence that a specific biological entity is an independent driver of prediction.

The same structure suggests a further, more speculative possibility: that graph-based models may be more robust to missing or corrupted inputs than models built on raw EHR features. Because each disease’s predictive signal is spread across many correlated features rather than concentrated in a few, the loss or corruption of any individual feature can be absorbed by the others; in the No-Graph models, where signal rests on a small number of near-independent features, the same perturbation would be expected to degrade performance more sharply. Our current results are consistent with this idea - graph-model performance was largely maintained as features were removed, whereas No-Graph performance declined under the same removal – but they probe feature removal and single-feature shuffling rather than realistic input noise, and so provide only indirect evidence. We therefore propose, as a hypothesis for future work, that knowledge-graph integration may confer resilience to the incompleteness and noise that characterize real-world EHR data, a form of value distinct from raw discriminative accuracy.

A further limitation is that our three cohorts differ simultaneously in biological specificity, in density of representation within SPOKE, and in whether the condition is a discrete disease at all (chronic pain being better regarded as a heterogeneous phenotype); because these properties covary, we cannot attribute the disease-dependent behavior to any single one of them. Our conclusions have a defined scope as they are drawn from three cohorts and a single knowledge graph (SPOKE), so whether integration utility can be predicted from disease characteristics remains to be tested at larger scale. Finally, our mechanistic account rests on correlation and ablation, which describes how predictive signal is organized across features rather than the independent information each feature contributes.

Our results open several additional future directions. The disease-dependent pattern observed here motivates evaluation across a broader range of cohorts spanning variation in biological specificity and representation within the knowledge graph, to determine whether the utility of integration can be anticipated from disease and graph characteristics and ultimately translated into an actionable design principle. Comparing diseases matched along some of these dimensions while differing along others, for example, two well-represented, genetically influenced neurodegenerative diseases such as Parkinson’s and Alzheimer’s disease, could help disentangle disease-intrinsic factors from differences in graph coverage and identify which characteristics determine when knowledge-graph integration is likely to be beneficial. Additionally, because our cohorts were built from administrative EHR coding, applying the approach to richer sources, such as intensive-care datasets like MIMIC that add laboratory values, vital signs, medication administration and clinical text, would establish whether diffusion contributes more when the underlying representation is itself richer, or whether it continues mainly to re-express existing signal. The robustness hypothesis raised by our feature analysis can be tested directly, by systematically removing or degrading portions of the input data and measuring how well graph-derived representations hold up compared with the raw EHR features. Alternative knowledge graphs, alternative diffusion parameterizations, and prospective or external validation would further map how far these effects extend. Pursuing these directions would determine how broadly the principles identified here - matching graph scope to the data, expecting disease-dependent returns, and treating the graph as a transformation of existing information, rather than a source of new signal - hold across diseases, data modalities and knowledge graphs.

## Methods

### SPOKE Knowledge graph

We used an early release of SPOKE (Scalable Precision Medicine Open Knowledge Engine)[4], obtained from the SPOKE project’s public repository. This release is substantially smaller than current versions of the graph, which have since expanded to millions of nodes; the version used here contains 47,031 nodes and 4,215,418 unique directed edges (Supplementary Fig. 1). The release stores 4,500,394 typed relationships, which collapse to 4,215,418 unique directed edges when relationship type is disregarded, as in the connectivity matrix used for PageRank propagation. SPOKE comprises 11 biological and clinical entity types: Gene (20,945), BiologicalProcess (11,381), SideEffect (5,734), MolecularFunction (2,884), Pathway (1,822), Compound (1,552), CellularComponent (1,391), Symptom (438), Anatomy (402), PharmacologicClass (345) and Disease (137). Edges encode typed, directed biological and clinical relationships, including gene– disease associations, drug–target interactions, pathway memberships and symptom–compound relationships (Supplementary Fig. 1). Notably, Disease nodes constitute only 0.3% of the graph (137/47,031), which directly constrains the size of disease-centered subgraphs and contextualizes the differences in biological neighborhood size across the three diseases studied.

### Patient Cohorts

All analyses used de-identified structured EHR data from UCSF Medical Center, comprising diagnosis codes (ICD-9 and ICD-10), medication orders, and laboratory results. Three disease cohorts: Parkinson’s Disease (PD), Multiple Sclerosis (MS), and Chronic Pain (CP), were constructed to represent a spectrum of biological specificity, clinical heterogeneity, and SPOKE knowledge graph representation.

### Cohort Matching

For all three disease cohorts, demographic variables were not available for matching, so we characterized each patient by their healthcare utilization and used this to construct balanced disease and control groups. Each patient’s healthcare footprint was defined as the number of distinct EHR concepts that mapped to SPOKE (equivalently, the number of active SPOKE Entry Point or *SEPs* nodes in their record). Cases and controls were then matched 1:1 on this footprint: proceeding through cases one at a time, each case was paired with the as-yet-unused control whose footprint was closest to it (greedy matching without replacement), yielding cohorts with comparable levels of recorded healthcare activity. Matching was performed prior to graph construction and feature-matrix ablation, so that all downstream analyses operated on cohorts balanced for clinical complexity. Across all three cohorts, every identified case mapped to SPOKE (none lost), and cases were matched from a shared pool of ∼145,000 control patients (146,543 patients × 3,183 SEP nodes in the global patient-by-SEP matrix).

#### Multiple Sclerosis (MS)

MS patients were identified using ICD-10 code G35. A total of 2,349 patients were identified and mapped to SPOKE. Following 1:1 EHR footprint score matching, the final cohort comprised 2,349 MS cases and 2,349 controls (n = 4,698).

#### Parkinson’s Disease (PD)

PD patients were identified using ICD-10 code G20 and ICD-9 code 332.0 (primary Parkinson’s disease). ICD-9 code 332.1 (secondary parkinsonism) was excluded to avoid including patients whose symptoms arose from medication side effects or other non-primary causes. A total of 2,149 PD patients were identified yielding a final cohort of 2,149 PD cases and 2,149 matched controls (n = 4,298 total).

#### Chronic Pain (CP)

Chronic Pain patients were identified using chronic-specific ICD10 G89 sub-codes (G89.0, G89.21, G89.22, G89.28, G89.29, G89.3, G89.4) and ICD9 chronic 338 sub-codes (338.0, 338.2, 338.28, 338.29, 338.3, 338.4); acute pain sub-codes were explicitly excluded (ICD10: G89.11, G89.12, G89.18; ICD9: 338.1x). In this cohort all 11,874 cases were captured by the chronic G89 codes (the 338 codes added no additional patients). The final cohort comprised 11,874 CP cases and 11,874 matched controls (n = 23,748). Chronic pain has no discrete Disease node in SPOKE: G89 is a symptom code, and chronic pain is represented only by two peripheral nodes-D059350 (a Symptom node) and C0150055 (a SideEffect node)-neither of which can be reached from patients’ records, because the G89/338 diagnosis codes do not resolve onto them through the standard ICD→CUI→SPOKE mapping. These two nodes have only 13 neighbors in SPOKE, too few to support meaningful diffusion, so a disease-centered subgraph could not be constructed for this cohort (see methods, *Disease-Graph construction*).

### Patient-SPOKE Mapping (SEP Matrix)

EHR concepts were mapped to SPOKE nodes using standardized ontologies: diagnosis codes (ICD-9/10) via UMLS Concept Unique Identifiers (CUIs) and Disease Ontology identifiers (to Disease and Symptom nodes); medication orders via DrugBank identifiers (to Compound nodes); and laboratory concepts via LOINC codes, which resolve through a CUI-based hierarchy onto existing SPOKE nodes. The resulting SPOKE Entry Points (SEPs)[4] are the subset of SPOKE nodes directly mappable to the EHR and serve as the biological entry points for graph propagation.

The SEPs were comprised of 3,183 nodes (1,851 SideEffect, 922 Compound, 257 Symptom, 137 Disease, 16 Gene). This mapping was performed once over the full patient population, producing a single binary patient × SEP matrix (146,543 × 3,183), where 1 indicates that a patient’s record contains a concept mapping to that node and 0 its absence.

For each disease, the matched cohort was obtained by extracting the corresponding rows from this global SEP matrix, yielding matched SEP matrices of 4,698 (MS), 4,298 (PD), and 23,748 (Chronic Pain) patients × 3,183 SEP nodes. Disease-associated leakage nodes were then removed to form the final feature sets (see, Methods-*Leakage Removal*). All downstream graph construction, PageRank propagation, and feature-matrix assembly used the cohort-specific matched SEP matrix as the starting point.

### Enrichment Analysis

To identify SPOKE nodes enriched in cases relative to matched controls, a one-tailed Fisher’s exact test was applied to each of the 3,183 SEP nodes in the matched cohort. Multiple-testing correction used the Benjamini–Hochberg false-discovery-rate procedure. Nodes were flagged as candidate leakage nodes if they met all three criteria: FDR q < 0.05, odds ratio > 3, and case prevalence > 5%. This node-level test was used specifically to identify candidate leakage features and was computed prior to leakage removal. It provided the empirical basis for identifying candidate leakage nodes. The complementary cohort-level view - characterizing how strongly and specifically each disease is represented across the EHR - is shown in Figure 1b.

### Leakage Removal

Enriched nodes were reviewed as candidate leakage features-nodes whose presence acts as a near-direct proxy for the disease label rather than a biologically informative association and evaluated against their prevalence among all available controls in the full unmatched cohort (Supplementary Table 2). Two types of leakage were removed: First, disease-defining concepts with near-zero control prevalence (≤ 0.20%): the disease diagnosis nodes themselves (100% of cases, 0% of controls) and disease-specific medications (fingolimod, modafinil and optic neuritis in MS; rasagiline, entacapone and L-DOPA in PD; each present in ≤ 0.14% of controls). Second, for chronic pain, symptom concepts that directly encode the pain phenotype (back pain, low back pain and loin pain); although common in controls (∼9-10%), these constitute the clinical definition of the disease and therefore act as label proxies. Chronic pain has no diagnosis node in SPOKE and no disease-specific medication, so no near-zero-prevalence leakage existed for this cohort.

To confirm that our findings were not sensitive to the stringency of leakage removal, we defined two feature sets per cohort (Supplementary Table 2). The final feature set, used for all analyses reported removed the complete leakage set - including the single most disease-defining feature in each cohort (L-DOPA in PD; optic neuritis in MS; the pain-concept nodes in CP), yielding 3,178 features per cohort. A less-stringent feature set, used for a sensitivity analysis, removed only the core leakage nodes (the diagnosis and disease-specific medications) but retained these most disease-defining markers. All conclusions were unchanged between the two schemes (Methods, *Sensitivity to leakage-removal stringency*; Supplementary Table 2, Supplementary Fig. 2)

### Graph Construction Strategies

Five strategies of increasing graph integration were evaluated per cohort:

***No-Graph:*** used the raw binary SEP features (3,178) with no graph integration, trained directly with a random forest- the standard machine-learning baseline against which all graph-based strategies were compared.

The remaining four generated patient embeddings by personalized PageRank diffusion (PSEV) and differed only in the graph over which diffusion was performed.

***Cohort-Graph***: This sub-graph was constructed by taking all SEP nodes observed in the matched cohort and expanding one hop into SPOKE, thus retaining all neighbors except hub nodes (degree > 500; 1,072 hubs removed), yielding ∼21,200-node subgraphs (PD 21,139; MS 21,204; CP 21,225).

***Concept-Graph:*** This graph was constructed using a minimal subgraph of the SEP nodes alone (3,178), retaining edges only where both endpoints were SEP nodes, with no expansion beyond the EHR-mapped concept set; this isolates the contribution of direct connections among observed EHR concepts.

***Disease-Graph:*** This sub-graph was expanded using one hop (all edge types, no hub removal) from disease seed nodes: PD: DOID:14330 and C0030567 (798 nodes); MS: DOID:2377 and C0026769 (322 nodes). For chronic pain, the Disease-Graph was excluded: its seed nodes (D059350, C0150055) are peripheral Symptom and SideEffect nodes with only 13 neighbors in SPOKE, too sparse to support meaningful diffusion.

***Whole-Graph:*** Computed PSEVs over the complete SPOKE graph (47,031 nodes, 4,215,418 edges) without subsampling, the maximum available biological context, reproducing the original SPOKEsig framework.

### Computational Parameters

All modelling strategies were evaluated under identical computational and analytical parameters, so that differences in runtime reflected graph scope alone. Computations were performed on a single server (Intel Xeon Silver 4214, 2.20 GHz; 48 logical cores; 188 GB RAM).

### PSEV Computation

Patient information was embedded onto SPOKE using Propagated SPOKE Entry Vectors (PSEVs), computed with a modified personalized PageRank algorithm following Nelson et al. A separate PSEV was computed for each SEP node. Rather than restarting to a single node, the random walker restarted-with probability 0.1, to a population-level co-occurrence prior: for every patient in the matched cohort, weight was distributed uniformly across the SEP nodes present in that patient’s record, and these contributions were aggregated across the cohort and normalized per SEP, so that each SEP’s restart distribution reflected how frequently the other EHR-mapped concepts co-occurred with it across the population. PageRank was iterated with a restart probability of 0.1, until the change in the rank vector between successive cycles fell below a tolerance of 0.001, up to a maximum of 40 iterations; the resulting vector assigns every node in the graph a score encoding its relevance to that entry point. Computations were parallelized across 48 workers (Python multiprocessing with shared memory) and run over the complete node set of each graph, with no subsampling. Convergence was tracked for every computation. Whereas the original PSEV implementation iterated to convergence without a fixed iteration limit, we capped iterations at 40 for tractability across the large node sets evaluated here. Under this cap, the majority of SEP-node PSEVs in the Cohort-Graph reached the iteration limit without meeting the tolerance (>99% across all three cohorts), whereas PSEVs over the Whole-Graph converged far more often (≈50% of SEP nodes in MS). This pattern was consistent across diseases and reflects the topology of the cohort-derived subgraph together with the iteration cap, rather than any disease-specific property.

### SPOKEsig Feature Construction

Patient-level SPOKEsig vectors were constructed by multiplying each patient’s binary SEP vector by the PSEV matrix, yielding a raw score for every node in the graph reflecting that node’s relevance to the patient’s EHR profile. Scores were rank-transformed row-wise per patient (ties assigned the minimum rank) and the resulting rank matrix was z-score normalized across all patients. Each strategy’s complete feature matrix combined the binary SEP features with the corresponding SPOKEsig scores, giving final-set matrices of 6,356 features (Concept-Graph), 24,317-24,403 features (Cohort-Graph, cohort-dependent) and 50,209 features (Whole-Graph); the No-Graph baseline used the 3,178 SEP features alone

### Classification and Evaluation

Disease classification was performed using Random Forest (500 trees, all available cores). No feature scaling was applied, as Random Forests are scale-invariant. Model performance was evaluated using the area under the receiver operating characteristic curve (AUROC). To obtain stable and mutually independent performance estimates, we used a 1×25 independent-build design: 25 independent random seeds were drawn from a master seed (0) using numpy, and for each seed a single stratified 80/20 train-test split was generated as fold 0 of a stratified five-fold partition (StratifiedKFold, shuffle = True, seeded by that seed). A Random Forest was trained and evaluated on each split, yielding 25 independent AUROC estimates per method. Unlike repeated cross-validation, these 25 builds share no common training data and therefore provide independent estimates. The Random Forest’s own random state was varied across builds (set to, and recorded with, each build’s seed), so that model stochasticity as well as the data partition varied across the 25 estimates. The same 25 seeds, and hence the same train-test splits, were used across all diseases and graph configurations, ensuring that identical test patients appeared across methods within each disease; a critical design choice enabling paired statistical comparison. Mean AUROC, standard deviation, and 95% confidence intervals (t-distribution, df = 24) were computed across the 25 estimates.

### Sensitivity to leakage-removal stringency

To confirm that findings did not depend on how aggressively leakage was removed, the full modelling pipeline was repeated on the supplementary feature set (core leakage removed only; the borderline markers of Supplementary Table 2 retained) and compared with the final, more conservative set used in the main analyses. As expected, retaining the additional disease-defining features modestly raised absolute AUROC (Supplementary Fig. 2), most in the cohorts whose retained feature was strongly predictive (PD, L-DOPA retained: No-Graph AUROC 0.961 vs 0.944; chronic pain, pain-concept nodes retained: 0.843 vs 0.818) and minimally in MS (optic neuritis retained: 0.942 vs 0.941). All qualitative conclusions were unchanged: the Cohort-Graph matched or exceeded the Whole-Graph in every cohort; the incremental value of graph integration over the No-Graph baseline remained positive for PD, small but positive for MS, and negative for chronic pain; and the Disease-Graph collapsed in both PD and MS. The final, more conservative feature set is therefore reported throughout, with the less-stringent set confirming robustness (Supplementary Fig. 2).

### Statistical Analysis

Pairwise differences in predictive performance between graph configurations were assessed by two-sided paired t-tests on the 25 AUROC estimates (scipy.stats.ttest_rel), with Bonferroni correction applied within each cohort (10 pairwise comparisons for MS and PD; 6 for chronic pain, which has no Disease-Graph). Estimates were paired by seed and fold, so that identical held-out patient sets were compared across methods. Effect sizes were quantified using Cohen’s d for paired samples- the mean of the paired differences divided by their standard deviation (d_z), interpreted as negligible (|d| < 0.2), small (0.2-0.5), medium (0.5–0.8) or large (|*d*| > 0.8); because the paired differences are highly consistent across folds, this metric can take large values.

### Permutation Feature Importance

To characterise the mechanistic effect of graph integration on model behaviour, permutation feature importance was computed for three configurations per cohort: No-Graph (Pure RF), Concept-Graph (Zero-Hop with intra-SEP edges), and Cohort-Graph. To restrict computation to informative features, Gini importance (averaged across the 25 independent builds) was first used to select stable candidates, features with mean Gini importance > 0.001 (No-Graph: 225-246; Concept-Graph: 78-183; Cohort-Graph: 35-147 features per cohort, decreasing with graph scope and lowest in chronic pain). Permutation importance was then computed with a custom implementation replicating scikit-learn’s *permutation_importance*: for each stable feature, its column in the held-out test matrix was randomly shuffled while all other features were left intact, and the resulting drop in AUROC was recorded. This was repeated three times per feature in each of the 25 builds (75 shuffles per feature in total), and final importance was defined as the mean AUROC drop across all 25 builds. For each feature the output recorded the SPOKE node ID, node name, node type, feature source (SEP binary, a raw EHR flag; or SPOKEsig, a graph-propagated score), and the mean and standard deviation of the AUROC drop.

### Feature Ablation

To determine whether predictive signal was concentrated in a few features or distributed across many, we performed a cumulative feature-ablation analysis on the recruited feature set of each configuration. Within each of the 25 independent builds, recruited features were removed cumulatively in one of two orders: most-important-first, ranked by the consensus permutation importance, or in random order as a control (an independent random permutation seeded per build). After each removal, a new Random Forest with identical parameters was trained on the remaining features and evaluated on the same held-out test split, yielding test AUROC as a function of the number of features removed. Curves show the mean and standard deviation across the 25 builds, plotted against the percentage of recruited features removed; the Cohort-Graph full-matrix (all-features) AUROC and the 0.5 chance level are shown for reference. A steep, importance-dependent decline indicates reliance on a few features carrying independent signal, whereas a curve that stays near the full-matrix AUROC until most features are removed, and that is insensitive to removal order, indicates signal distributed across many correlated features.

### Feature Correlation Analysis

To characterise the correlation structure underlying these ablation profiles, we computed pairwise Pearson correlations among the recruited features of each configuration. For each cohort and graph scope, the top 20 recruited features (ranked by permutation importance) were selected, and the correlation matrix of their values across all patients was computed. Features were ordered within each matrix by hierarchical clustering (average linkage on a distance of 1 -|r|) to reveal blocks of correlated features, and each configuration was summarised by the median absolute pairwise correlation (median |r|).

## References

1. Jensen, P.B., L.J. Jensen, and S. Brunak, Mining electronic health records: towards better research applications and clinical care. Nat Rev Genet, 2012. 13(6): p. 395–405.

2. Rajkomar, A., et al., Scalable and accurate deep learning with electronic health records. NPJ Digit Med, 2018. 1: p. 18.

3. Xiao, C., E. Choi, and J. Sun, Opportunities and challenges in developing deep learning models using electronic health records data: a systematic review. J Am Med Inform Assoc, 2018. 25(10): p. 1419–1428.

4. Morris, J.H., et al., The scalable precision medicine open knowledge engine (SPOKE): a massive knowledge graph of biomedical information. Bioinformatics, 2023. 39(2).

5. Himmelstein, D.S., et al., Systematic integration of biomedical knowledge prioritizes drugs for repurposing. Elife, 2017. 6.

6. Chandak, P., K. Huang, and M. Zitnik, Building a knowledge graph to enable precision medicine. Sci Data, 2023. 10(1): p. 67.

7. Cowen, L., et al., Network propagation: a universal amplifier of genetic associations. Nat Rev Genet, 2017. 18(9): p. 551–562.

8. Nelson, C.A., et al., Embedding electronic health records onto a knowledge network recognizes prodromal features of multiple sclerosis and predicts diagnosis. J Am Med Inform Assoc, 2022. 29(3): p. 424–434.

9. Soman, K., et al., Early detection of Parkinson’s disease through enriching the electronic health record using a biomedical knowledge graph. Front Med (Lausanne), 2023. 10: p. 1081087.

10. Nelson, C.A., A.J. Butte, and S.E. Baranzini, Integrating biomedical research and electronic health records to create knowledge-based biologically meaningful machine-readable embeddings. Nat Commun, 2019. 10(1): p. 3045.

11. Errica, F., et al., A Fair Comparison of Graph Neural Networks for Graph Classification. 2020.

12. Shchur, O., et al., Pitfalls of Graph Neural Network Evaluation. CoRR, 2018. abs/1811.05868.

13. Bonner, S., et al., Implications of topological imbalance for representation learning on biomedical knowledge graphs. Briefings in Bioinformatics, 2022. 23(5).

14. Bonner, S., et al., A review of biomedical datasets relating to drug discovery: a knowledge graph perspective. Briefings in Bioinformatics, 2022. 23(6).

15. Strobl, C., et al., Conditional variable importance for random forests. BMC Bioinformatics, 2008. 9(1): p. 307.

16. Toloşi, L. and T. Lengauer (2011). "Classification with correlated features: unreliability of feature ranking and solutions." Bioinformatics 27(14): 1986–1994.

