## Supplementary_Material for "Understanding when and how knowledge graphs add value to EHR-based clinical prediction across Parkinson’s disease, multiple sclerosis, and chronic pain"

##### Supplementary Figure 1

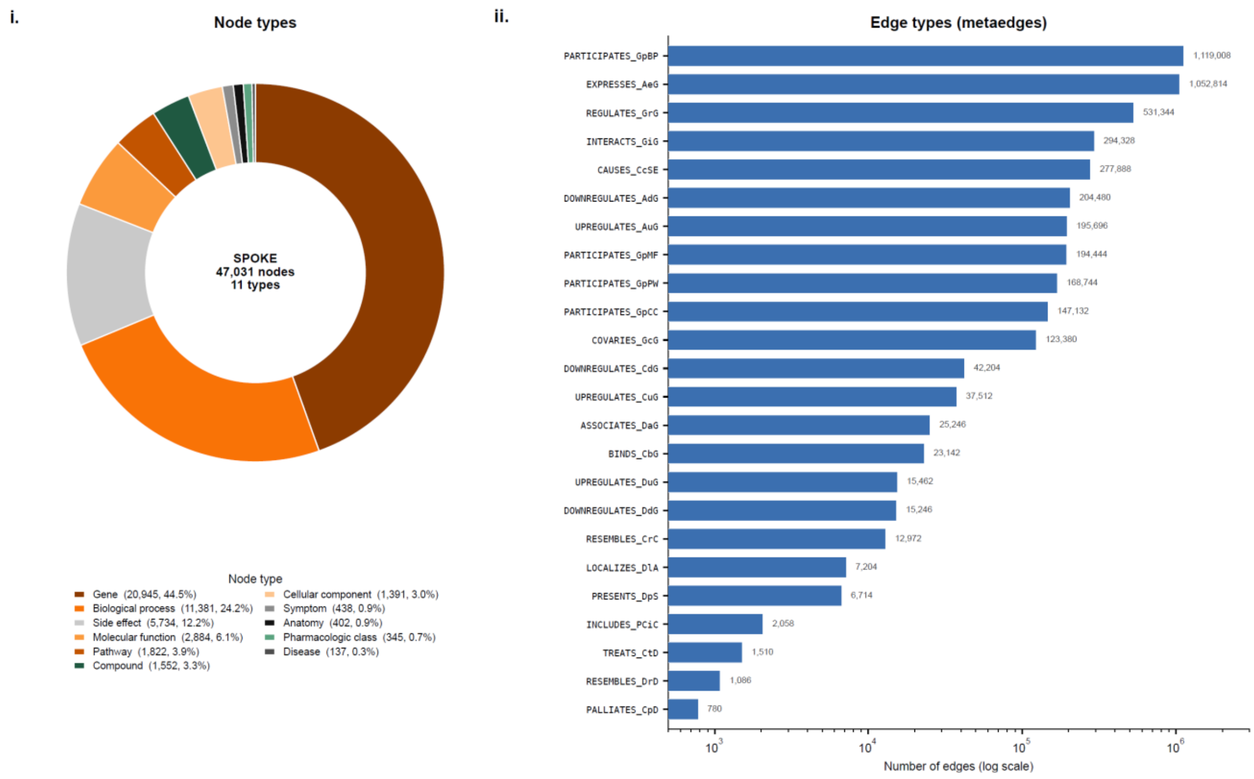


***Supplementary Figure 1. Composition of the SPOKE knowledge graph [4].*** *(i) Distribution of the 47,031 SPOKE nodes across 11 node types, dominated by genes (44.5%) and biological processes (24.2%). (ii) The 4,500,394 typed edges across the 24 SPOKE metaedge types (horizontal bars, log scale; counts labelled), spanning more than three orders of magnitude- from PARTICIPATES_GpBP (Gene-participates-Biological Process; 1,119,008) to PALLIATES_CpD (780); abbreviations follow SPOKE metaedge notation. Note that this total counts every typed edge in the graph; the personalized-PageRank diffusion used throughout our analyses operates on the connectivity matrix, in which parallel edges between the same node pair collapse to a single connection, yielding 4,215,418 unique node-pair connections - the ~4.2 million edges referred to in the main text and Figures 1-2.*

#### Supplementary Figure 2

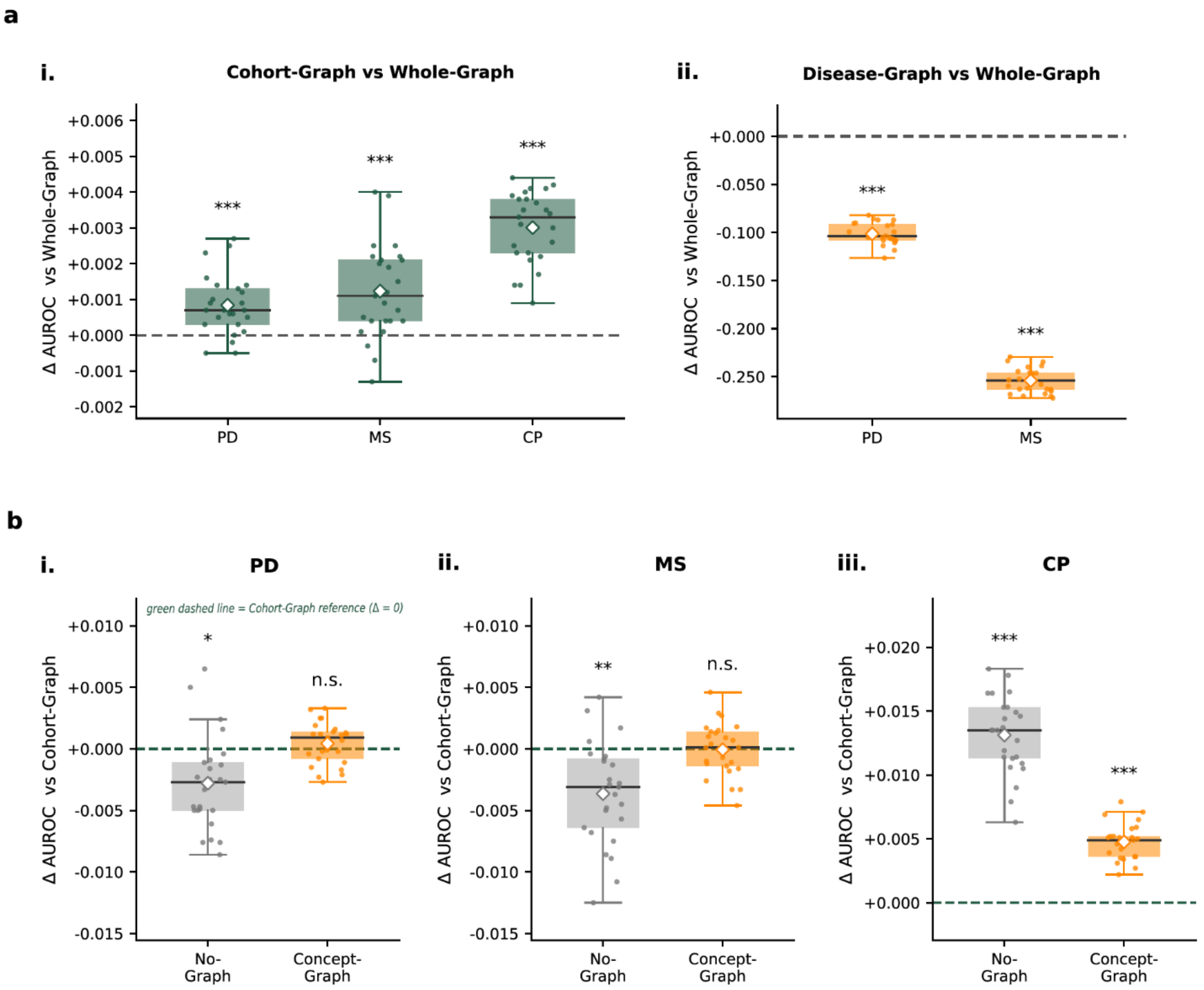


***Supplementary Figure 2. Sensitivity to leakage-removal stringency: main findings are preserved under the less-stringent feature set.*** *All analyses were repeated on the less-stringent feature set (core leakage removed only; the borderline markers of Supplementary Table 3 - optic neuritis [MS], L-DOPA [PD] and the back-pain concepts [CP] - retained), shown as per-instance AUROC differences across 25 independent build instances (two-tailed paired t-tests, Bonferroni-corrected; ** q < 0.01, *** q < 0.001; n.s., not significant). (a) Diffusion-strategy comparisons relative to Whole-Graph (mirroring Fig. 2a): (i) Cohort-Graph − Whole-Graph clusters near zero in every cohort (|Δ| < 0.005), and (ii) Disease-Graph − Whole-Graph collapses in PD and MS. (b) No-Graph and Concept-Graph relative to the Cohort-Graph reference (green dashed line, Δ = 0), per cohort (mirroring Fig. 3): in PD and MS the No-Graph baseline falls below the Cohort-Graph while the Concept-Graph is statistically indistinguishable from it, whereas in CP both exceed it. Effect directions match the primary analysis in all cohorts; the main quantitative change is a smaller No-Graph deficit in PD (significant at * rather than ***), because retaining the strongly predictive L-DOPA feature raises the graph-free baseline*

#### Supplementary Table 1

| Cohort | Inclusion (ICD-10) | Inclusion (ICD-9) | Exclusion | Cases (N) |
|---|---|---|---|---|
| MS | G35 (multiple sclerosis) | (none; identified by G35 only) | none | 2,349 |
| PD | G20 (Parkinson disease) | 332.0 (primary PD) | 332.1 (secondary parkinsonism), unless also G20/332.0 | 2,149 |
| CP | G89.0, G89.21, G89.22, G89.28, G89.29, G89.3, G89.4 | 338.0, 338.2, 338.28, 338.29, 338.3, 338.4 | acute-pain: G89.11/.12/.18; 338.1/.11/.12/.18/.19 | 11,874 |

***Supplementary Table 1. Cohort definitions.*** *ICD-9/ICD-10 inclusion and exclusion codes used to identify each disease cohort from UCSF structured EHR, with the resulting number of cases (each matched 1:1 to a control on EHR footprint). PD cases were captured by ICD-10 G20 (2,145) plus ICD-9 332.0 (4 additional); secondary parkinsonism (332.1) was excluded. Chronic pain used chronic-specific G89/338 sub-codes with acute-pain sub-codes excluded; all 11,874 CP cases were captured by the G89 codes (the 338 codes added none).*

### Supplementary Table 2

| Cohort | Concept name | SPOKE node | Node type | Cases, n (prevalence) | Controls, n (prevalence) | Enrichment (odds ratio) | FDR q-value |
|---|---|---|---|---|---|---|---|
| MS | multiple sclerosis | DOID:2377 | Disease | 2349 (100.00%) | 0 (0.00%) | ∞ (absent in controls) | <1e-300 |
| MS | multiple sclerosis | C0026769 | SideEffect | 2349 (100.00%) | 0 (0.00%) | ∞ (absent in controls) | <1e-300 |
| MS | Fingolimod | DB08868 | Compound | 192 (8.17%) | 0 (0.00%) | ∞ (Haldane ≈ 419) | 2.16e-57 |
| MS | Optic neuritis* | C0029134 | SideEffect | 166 (7.07%) | 0 (0.00%) | ∞ (Haldane ≈ 358) | 3.31e-49 |
| MS | Modafinil | DB00745 | Compound | 142 (6.05%) | 1 (0.04%) | 151.1 | 6.72e-40 |
| PD | Parkinson's disease | DOID:14330 | Disease | 2149 (100.00%) | 0 (0.00%) | ∞ (absent in controls) | <1e-300 |
| PD | Parkinson's disease | C0030567 | SideEffect | 2149 (100.00%) | 0 (0.00%) | ∞ (absent in controls) | <1e-300 |
| PD | L-DOPA* | DB01235 | Compound | 1189 (55.33%) | 3 (0.14%) | 886.0 | <1e-300 |
| PD | Rasagiline | DB01367 | Compound | 257 (11.96%) | 0 (0.00%) | ∞ (Haldane ≈ 585) | 4.99e-79 |
| PD | Entacapone | DB00494 | Compound | 229 (10.66%) | 0 (0.00%) | ∞ (Haldane ≈ 514) | 6.02e-70 |
| CP | Low back pain* | C0024031 | SideEffect | 4671 (39.34%) | 1063 (8.95%) | 6.6 | <1e-300 |
| CP | Low Back Pain* | D017116 | Symptom | 4671 (39.34%) | 1063 (8.95%) | 6.6 | <1e-300 |
| CP | Loin pain* | C0235632 | SideEffect | 4635 (39.03%) | 1060 (8.93%) | 6.5 | <1e-300 |
| CP | Back Pain* | D001416 | Symptom | 4364 (36.75%) | 1158 (9.75%) | 5.4 | <1e-300 |
| CP | Back pain* | C0004604 | SideEffect | 4364 (36.75%) | 1158 (9.75%) | 5.4 | <1e-300 |

***Supplementary Table 2. Disease-defining features removed prior to knowledge-graph analyses (leakage-removal).*** *All listed features were removed prior to modeling to prevent label leakage. Asterisk (*) marks features removed only in the primary analysis and retained in the less-stringent supplementary analysis (refer to Methods, Sensitivity to leakage-removal stringency) - for chronic pain every removed feature is asterisked, i.e. no features were removed under the less-stringent scheme. Odds ratio is cases versus matched controls (Fisher's exact test, FDR-corrected q); ∞ denotes zero prevalence in controls (odds ratio undefined), with the Haldane-Anscombe corrected estimate (+0.5 per cell) shown in parentheses where informative.*

Confidentiality Level: GENERAL/INTERNAL
